# Endometriotic Organoids: A Novel In Vitro Model of Endometriotic Lesion Development

**DOI:** 10.1101/2022.02.15.480583

**Authors:** Yong Song, Gregory W. Burns, Niraj R. Joshi, Ripla Arora, J. Julie Kim, Asgerally T. Fazleabas

## Abstract

The development and progression of endometriotic lesions are poorly understood, but immune cell dysfunction and inflammation are closely associated with the pathophysiology of endometriosis. A lack of suitable 3D *in vitro* models permitting the study of interactions between cell types and the microenvironment is a contributing factor. To address this limitation, we developed endometriotic organoids (EO) to explore the role of epithelial-stromal interactions and model peritoneal cell invasion associated with lesion development. Using a non-adherent microwell culture system, spherical organoids were generated with endometriotic epithelial cells (12Z) combined with immortalized endometriotic stromal cells (iEc-ESC) or immortalized uterine stromal cells (iHUF). Organoids self-organized with stromal cells occupying the center and epithelial cells on the periphery of the organoid. Endometriotic organoids (EO), containing iEc-ESC, resulted in the development of stratified 12Z epithelial cells compared to those with iHUF where the 12Z cells developed as a single layered epithelium. Transcriptomic analysis found 4,522 differentially expressed genes (DEG) between EO and 12Z/iHUF organoids, and the top DEG included increased expression of interleukins and prostaglandin synthase enzymes. An overlap of the EO DEG with baboon endometriotic lesions was highly significant. Finally, to mimic invasion of endometrial tissue into the peritoneum, a model was developed using EO and extracellular matrix containing human peritoneal mesothelial cells (LP9). Invasion of EO into the extracellular matrix-LP9 layer was increased in presence of estrogen or THP1-derived proinflammatory macrophages. Taken together, our results strongly support the concept that EO are an appropriate model for dissecting mechanisms that contribute to endometriotic lesion development.

**One Sentence Summary:** Endometriotic organoids are an appropriate model to study epithelial-stromal interactions and model cell invasion associated with lesion development.

## INTRODUCTION

Endometriosis is a chronic inflammatory hormone-dependent disorder which is characterized by the presence of endometrial tissue outside the uterus (*1*). It affects between 10 and 15% of all women of reproductive age, 70% of women with chronic pelvic pain, and 20–50% women with infertility (*2, 3*). An average 7-10 years delay between the onset of symptoms and definitive diagnosis places an enormous emotional and financial burden on both the patient and the healthcare system, with annual costs exceeding $78 billion each year in the United States alone (endofound.org). Quality of life is reduced in women with endometriosis due to painful symptoms, which result in a loss of 11 hours of productivity per week (endofound.org). The etiology and pathogenesis of endometriosis remains unclear, but lesion development has been associated with cellular proliferation and invasion, and interactions between endometriotic cells, immune cells and their microenvironment (*4, 5*). Appropriate *in vitro* platforms that allow dissection of the underlying mechanisms of these processes will inform the understanding of the disease and development of novel therapeutics.

Clinical samples from patients with endometriosis are the first line model to investigate the disease. One of the major limitations to understanding the pathological events associated with endometriosis is the significant delay of 8-11 years for diagnosis and subsequent variations in disease etiology and progression. Thus, both *in vivo* and *in vitro* models are indispensable to study the cellular and molecular mechanisms associated with endometriotic lesion development at different stages, The non-human primate and rodent models of endometriosis have been used most exclusively for *in vivo* studies. Non-human primates offer a physiologically relevant model and provide an opportunity to study both the early events and the disease progression (*6, 7*). Rodent models are advantageous due to the ability for tissue-specific gene manipulation and initial *in vivo* therapeutic testing (*8*). Most *in vitro* studies utilize two-dimensional monolayer culture of primary or immortalized endometrial cells. This method allows for the study of endometriosis associated mechanisms in a cell-specific manner and is easily manipulated for testing of treatments. However, the interactions are limited to the horizontal plane and cells are exposed to a uniform concentration of factors, so monolayer cultures are unable to mimic the architecture and function of tissues *in situ*. Thus, there is a significant gap between *in vivo* models and current *in vitro* cell culture techniques. Three-dimensional (3D) spheroids, or organoids, provide an exciting method to bridge the gap by incorporation of more complex structures and multiple cell types.

Organoids are self-organizing, genetically stable, 3D culture systems containing both progenitor/stem and differentiated cells that resemble the tissue of origin (*9-13*). Boretto et al. first established endometriotic epithelial organoids in a Matrigel organoid culture system using ectopic lesions from patients with endometriosis and reported that the organoids replicated disease-associated functional and genomic traits (*14*). This system only allowed for the culture of endometriotic epithelial cells. However, endometrial assembloids containing epithelium and stroma were possible when epithelial organoids were later combined with stromal cells in a collagen hydrogel (*15*). Wiwatpanit et al. generated endometrial organoids containing both epithelial and stromal cells from endometrial biopsies using micro-molded agarose plates and the scaffold-free organoids resembled the physiology of the normal endometrium (*13*). The micro-molded agarose plate organoid culture system is more cost effective, does not require exogenous scaffold materials but relies on *de novo* secretion of extracellular matrix, and allows for self-organization and growth of multiple cell types.

An SV40-tranformed endometriotic epithelial cell line derived from peritoneal endometriosis, 12Z, is widely used for the study of endometriosis (*16*) and we recently developed an endometriotic stromal cell line (iEc-ESC) from ovarian endometriosis (*17*). In this study, our aim was to develop an endometriotic organoid model containing both endometriotic epithelial and stromal cell lines. We hypothesized that the transcriptome of scaffold-free organoids containing both endometriotic cell types would mirror endometriotic lesions from the baboon model and provide a model to explore peritoneal invasion associated with endometriotic lesion initiation.

## RESULTS

### Epithelial and Stromal Cells Self-Organize in Organoids

Epithelial and stromal cells are the two main cell types found in endometriotic lesions that are hypothesized to be descended from retrograde menstrual endometrial fragments. The cells employed in this study were immortalized ectopic epithelial cells (12Z), ectopic stromal cells (iEc-ESC) and human uterine fibroblasts (iHUF) (*16, 17*). For live visualization of cell-type organization, epithelial 12Z cells were modified to express red fluorescent protein (RFP) and both stromal cell lines to express Azurite blue. Endometrial organoids were generated from epithelial 12Z or stromal iEc-ESC alone, 12Z+iHUF, and 12Z+iEc-ESC in micro-molded agarose well plates. A preliminary study empirically determined that the optimal ratio of epithelial cells to stromal cells for co-culture was 1:50. Time-lapse imaging demonstrated that epithelial and stromal cells reorganized autonomously in the agarose wells (Supplemental Video 1). The organization was confirmed by confocal image Z-stack reconstructions and immunohistochemical staining on days 4 and 7 (Fig 1). In 12Z+iHUF (Fig 1A) and 12Z+iEc-ESC (EO) (Fig 1B) organoids, epithelial cells were located on the periphery of a stromal cell core by day 4. EO developed multiple epithelial cell layers (Fig 1B), while 12Z+iHUF resulted in a single layer of epithelial cells at day 7 (Fig 1A). Organoids containing only epithelial 12Z or stromal iEc-ESC formed simple spheroid structures (Supplemental Fig 1).

**Fig 1.**
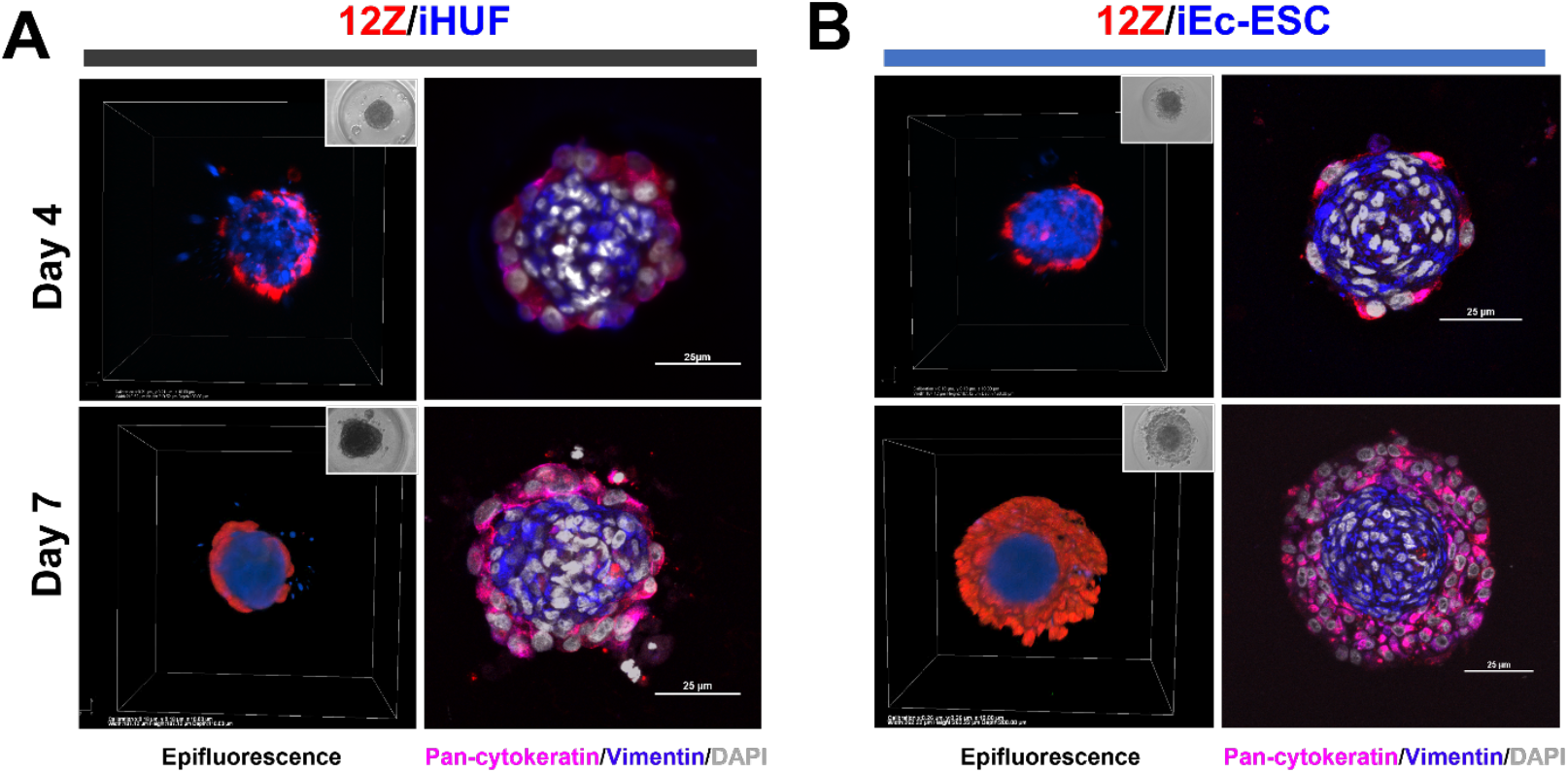
Organoid structure. Time course of organoid development derived from (A) 12Z-RFP/iHUF-Azurite Blue and (B) 12Z-RFP/iEc-ESC-Azurite Blue (endometriotic organoids, EO). The left panels are 3D views and inserts are phase contrast images. The right panels are immunofluorescent staining of vimentin (blue) and pan-cytokeratin (red). Scale bar = 25 μm.

### Endometriotic Organoids Invade a 3D Model of the Peritoneum

Cell invasion is one of the key processes involved in endometriotic lesion initiation and development. To simulate cell invasion that occurs *in vivo*, we developed a 3D endometriotic organoid invasion model. As shown in Fig 2A, human peritoneal mesothelial cells, LP9, expressing GFP were grown in Matrigel to mimic the structure of the peritoneum. Endometriotic organoids containing iEc-ESC and 12Z were seeded on the top of the LP9/Matrigel layer. The invasion of organoids was tracked using a fluorescent confocal microscopy every other day for 7 days. The Z-stack images were reconstructed to produce 3D models with Imaris image analysis software by Bitplane. As shown in Supplemental Video 2 and Fig 2C, the organoids invaded the Matrigel and continued penetrating the LP9 cell layer. Interestingly, stromal cells were always noted at the leading edge of invasion and as the first cell type to penetrate the LP9 cell layer. The organoid invasion was quantified, and data are shown in Fig 2B. Combined, these results support the idea that the organoid invasion model could be an ideal *in vitro* tool to study the early events of endometriosis, particularly with respect to cell invasion.

**Fig 2.**
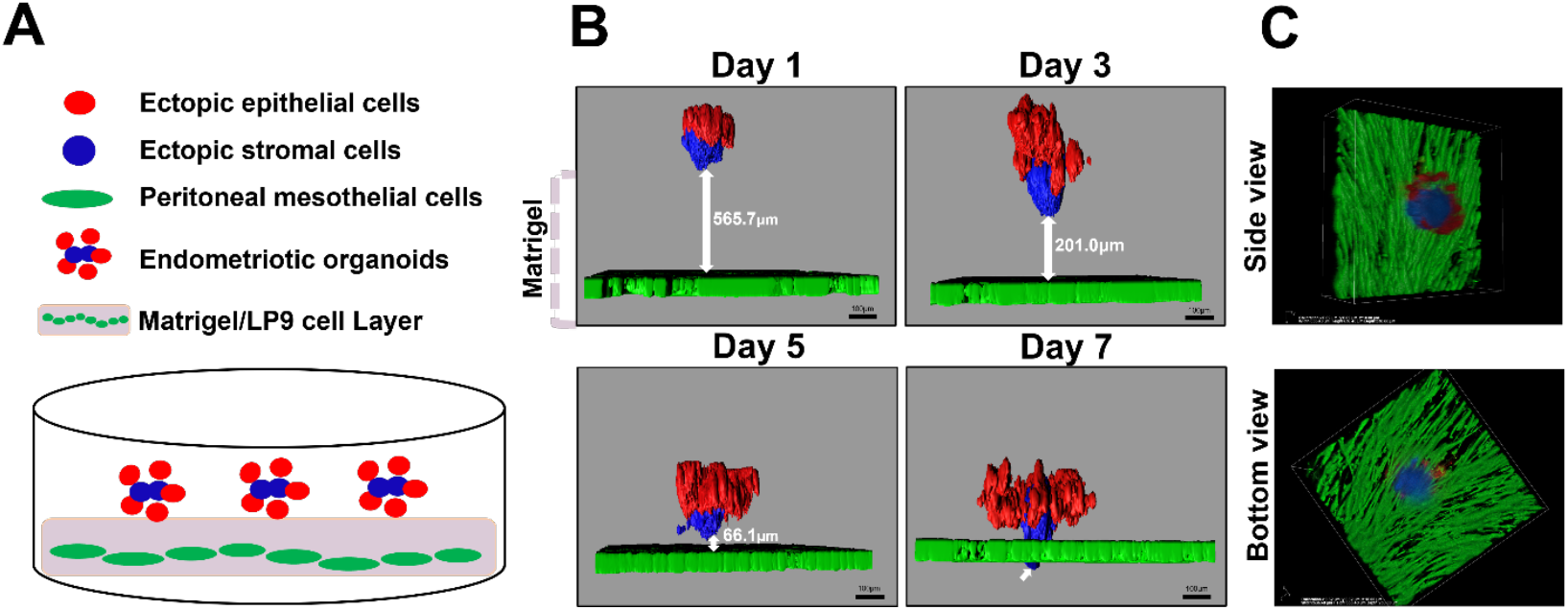
Organoid invasion model. Endometriotic organoids invade a 3D model of the peritoneum. (A) Schematic of the organoid invasion model. (B) Time course of organoid invasion quantified with Imaris image analysis software and relative to invasion into the mesothelial cell layer. Ectopic stromal cells were the leading edge of invasion through the LP9 cells. Note the penetration of organoids through the extracellular matrix and mesothelial cell layer in a time dependent manner. (C) Representative side view and bottom view of an EO invading the LP9 cells with stromal cells at the leading edge.

### Estrogen Stimulates Invasion of Endometriotic Organoids

Endometriosis is estrogen-dependent disease and estrogen can promote the migration and invasion of endometrial stromal cells and endometriotic epithelial cells in two-dimensional culture (*18, 19*). However, the effect of estrogen on organoids is unknown but we hypothesized that estradiol would stimulate invasion in our model. Invasion of Day 4 EO through the Matrigel/LP9 cell layer was monitored in the presence of 20nM estradiol (E2) or vehicle for 4 days. Organoids were imaged on Day 1 and Day 4 by fluorescent confocal microscopy. The invasion rate of endometriotic organoids was increased at 1.8-fold in the presence of E2 compared to vehicle (p<0.05; Fig 3).

**Fig 3.**
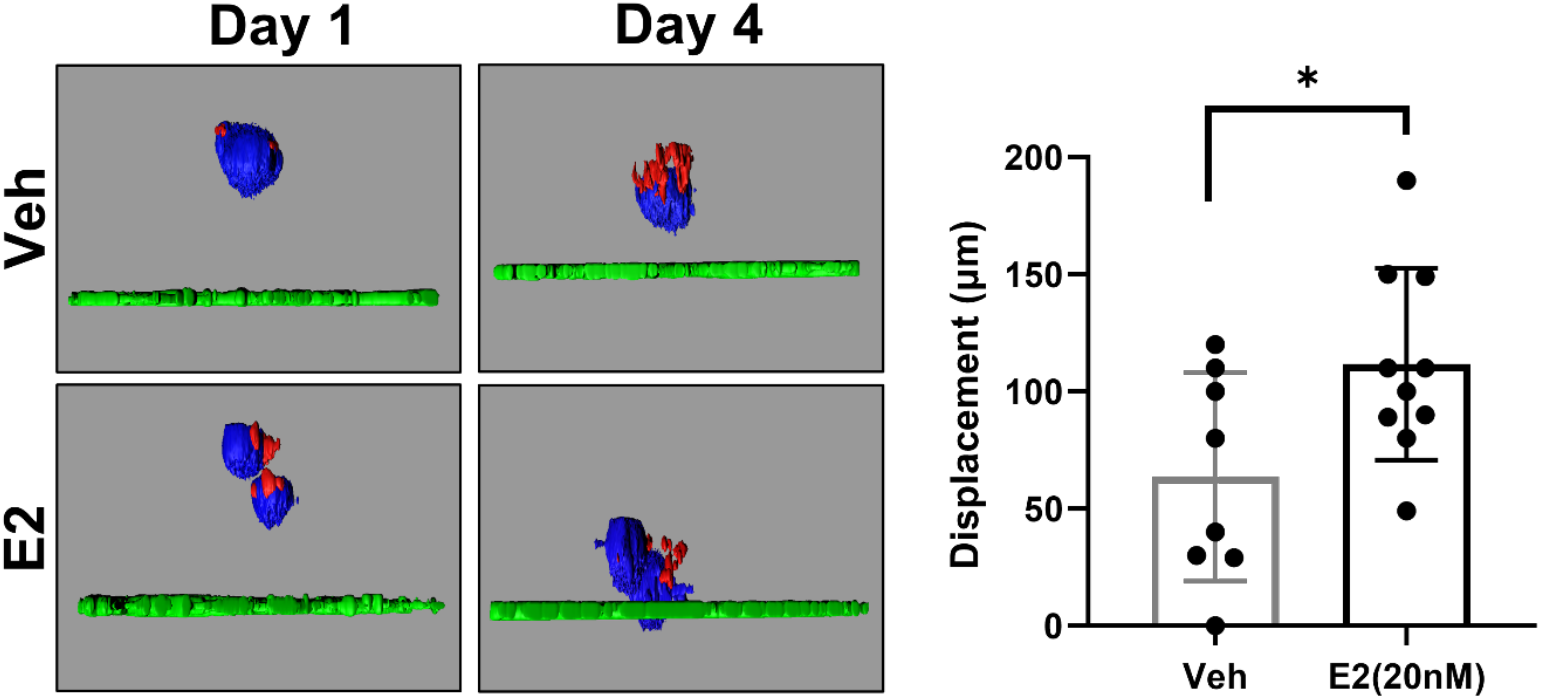
Estrogen stimulates invasion of endometriotic organoids (EO). Day 4 EO were seeded on an LP9/Matrigel layer and treated with vehicle (ethanol, n=8) or estradiol (E2, 20nM, n=10) for 4 days. The EO were imaged on days 1 and 4 using fluorescent confocal microscopy and invasion, relative to the LP9 cell layer was quantified. Invasion was increased by 1.8-fold in the presence of E2. *P<0.05

### Stromal Cell Origin Alters Organoid Gene Expression

We profiled the transcriptomes by RNA-sequencing of organoids containing ectopic stromal and epithelial cell lines (iEc-ESC/12Z; EO) and compared the profile to normal stromal and ectopic epithelial organoids (iHUF/12Z) to identify the global transcriptomic effect of ectopic stromal cells in the organoids. As an indication of broad transcriptome differences, EO and iHUF/12Z organoid samples were separated by principal component analysis across PC1 (65.7%, Fig 4A) and unsupervised hierarchical clustering (Fig 4B). Indeed, there were 4,522 differentially expressed genes (DEG), with 2,175 increased and 2,347 decreased, in EO compared to iHUF/12Z organoids. The top 50 DEG included increased expression of interleukins (*IL1B, IL24*), prostaglandin synthase enzymes (*PTGS1, PTGS2*), and telomerase reverse transcriptase (*TERT*). Of note, stromal cell lines were immortalized by constitutive expression of *TERT* via lentiviral transduction, but *TERT* was not differentially expressed in an analysis of organoids containing only stromal cells (Supplemental Fig 2), indicating the altered expression was due to an interaction between the epithelial and stromal cells.

**Fig 4.**
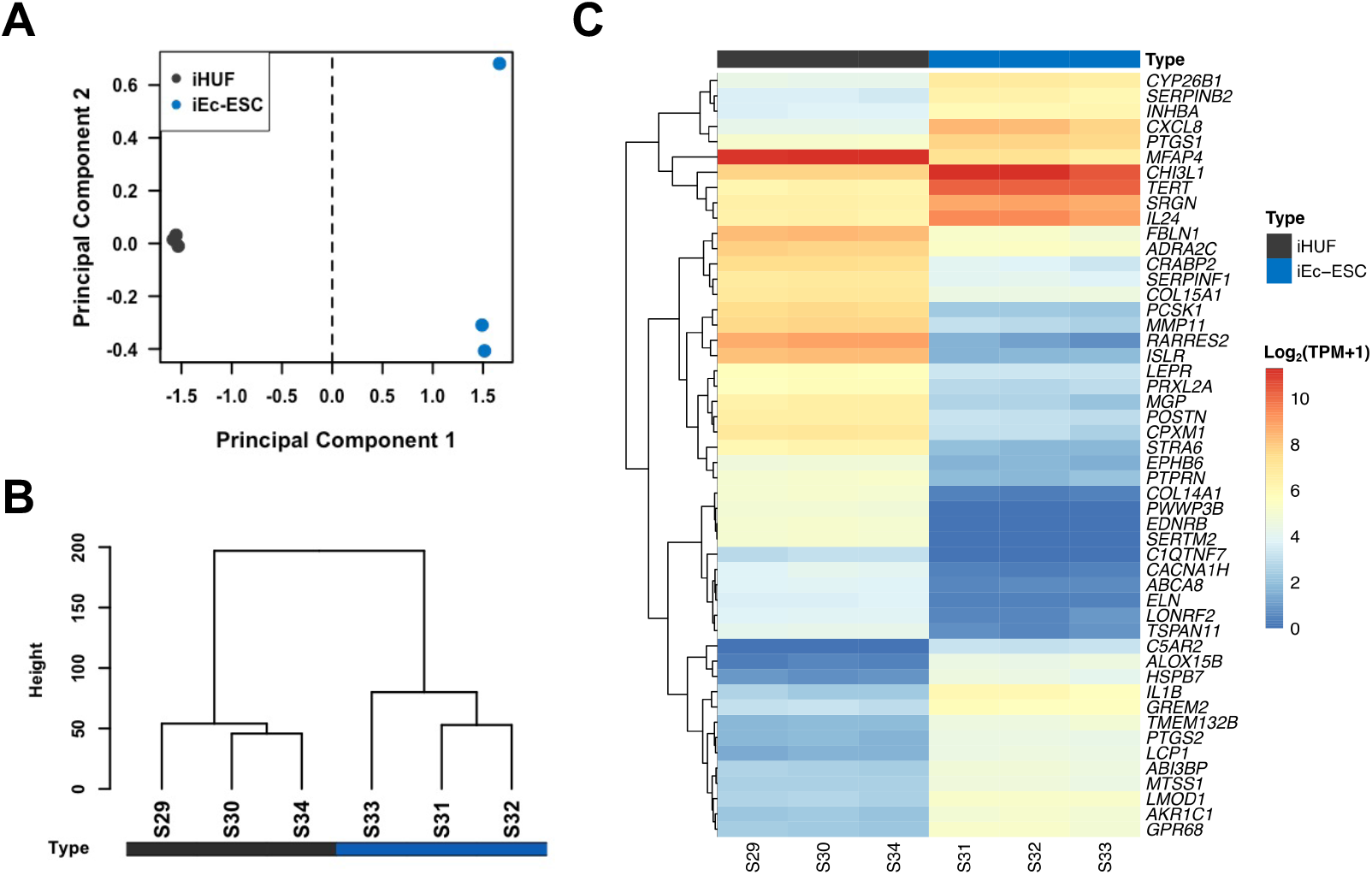
Stromal cell origin alters organoid gene expression. (A) Principal component plot demonstrating transcriptome separation between iEc-ESC/12Z (EO) and iHUF/12Z organoids. (B) Unsupervised hierarchical clustering dendrogram confirming separation of sample groups and (C) a heat map of the top 50 differentially expressed genes with hierarchical gene clustering.

Prostaglandin synthase enzymes, *PTGS1* and *PTGS2, CXCL8, TIMP3*, an inactivator of extracellular matrix remodeling enzymes, *IL1A*, and *LIF*, an estrogen response gene, were increased in EO while *STRA6* and *CRABP2*, retinoic acid binding proteins, and *PGR* were down-regulated (Fig 5A). Gene set enrichment analysis (GSEA) results included NFKB target genes, two senescence gene sets, and TNFα response via p38 mitogen-activated protein kinases as the most highly increased gene sets for EO, while stem cell-associated genes, including *CDKN2C, CDH11*, and *ROR1*, were decreased (Fig 5B).

**Fig 5.**
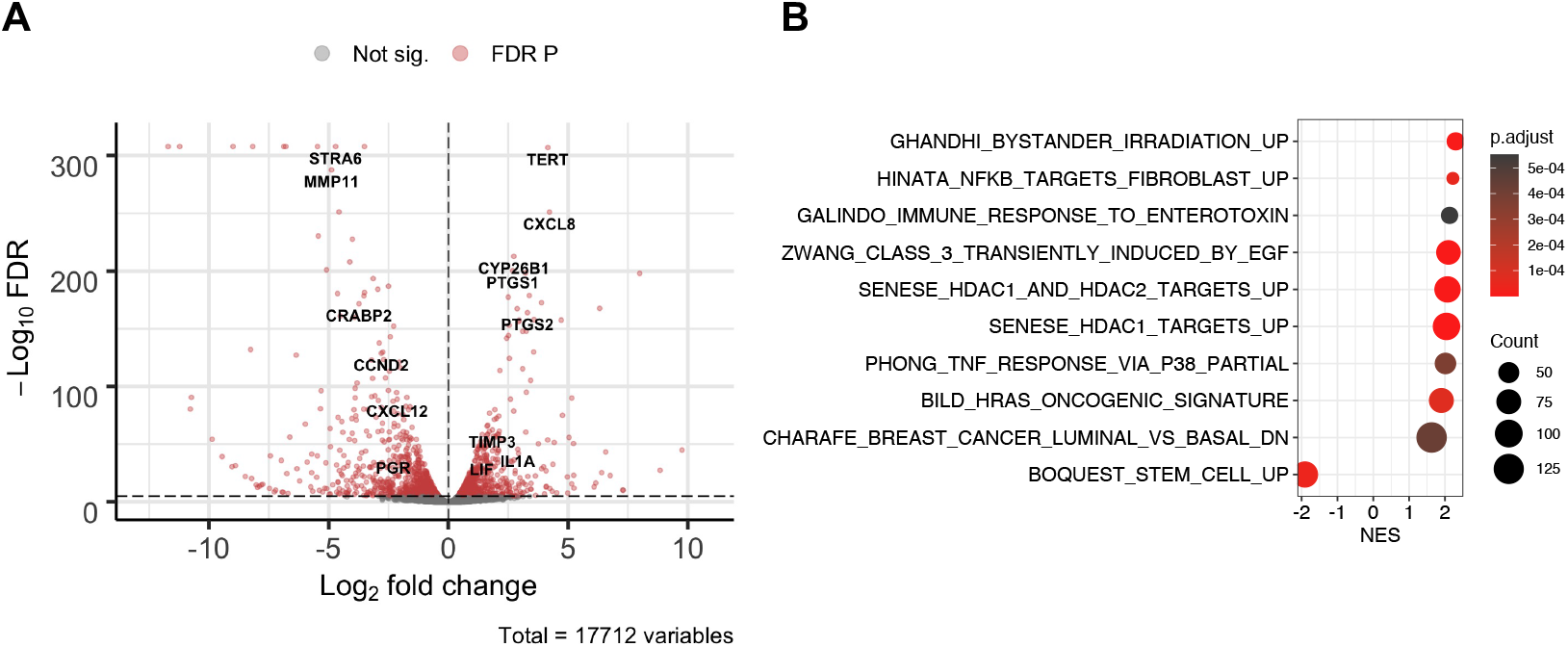
Transcriptome alterations in EO versus iHUF/12Z organoids. (A) Volcano plot highlighting several top downregulated and upregulated genes in EO. (B) The top MSigDB C2 enriched gene sets from human organoids. A positive normalized enrichment score (NES) indicates activation, or increased expression, of the gene set in EO.

### Ectopic Organoids are Similar to Endometriotic Lesions

RNA-sequencing data from baboon endometrium and endometriotic lesions was compared to the human organoids and overlap of gene expression between these comparisons was highly significant, with 12,504 commonly expressed genes (Fig 6A). Additionally, GSEA using hallmark gene sets found similarities for comparisons of EO versus iHUF/12Z organoids and spontaneous baboon ectopic lesions versus eutopic endometrium (Fig 6B). TNF signaling via NFKB, inflammatory responses, and hypoxia associated genes were increased in EO and baboon lesions, while interferon alpha response was decreased in EO and increased in lesions. Similarly, allograft rejection and IL6 JAK/STAT3 signaling gene sets were increased in baboon ectopic lesions compared to eutopic tissues but absent from the organoid comparison. Gene ontology analysis of 2,518 genes identified only as expressed in baboon tissues resulted in significant enrichment for PANTHER GO-slim terms associated with the immune system (Table 1).

**Fig 6.**
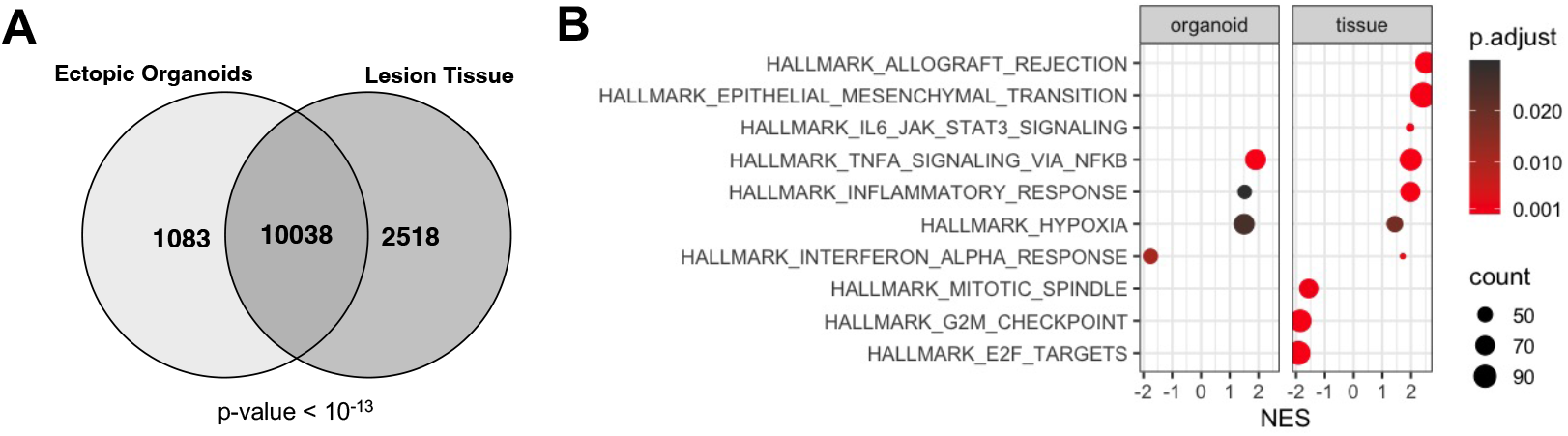
Endometriotic organoids (EO) and baboon lesion gene expression signatures are similar. (A) The overlap of all expressed genes in EO and baboon lesions from spontaneous endometriosis is highly significant. (B) Gene set enrichment analyses showed similar enrichment of NFKB, inflammatory response and hypoxia genes. NES= normalized enrichment score

**Table 1.** Enrichment for immune-related genes in spontaneous endometriotic lesions. Top 10 enriched biological process gene ontology terms from genes expressed in baboon lesions and not human organoids.

| PANTHER GO Slim (BP) | REFLIST | Input | Expected | Direction | Fold Enrichment | p-value | FDR |
| --- | --- | --- | --- | --- | --- | --- | --- |
| immune system process (GO:0002376) | 258 | 128 | 44.67 | + | 2.87 | 3.56E-20 | 7.74E-18 |
| immune response (GO:0006955) | 163 | 90 | 28.22 | + | 3.19 | 9.35E-17 | 1.13E-14 |
| cell surface receptor signaling pathway (GO:0007166) | 544 | 177 | 94.18 | + | 1.88 | 1.44E-12 | 1.49E-10 |
| signaling (GO:0023052) | 1208 | 321 | 209.14 | + | 1.53 | 2.95E-12 | 2.92E-10 |
| cell communication (GO:0007154) | 1219 | 320 | 211.05 | + | 1.52 | 1.29E-11 | 1.08E-09 |
| multicellular organismal process (GO:0032501) | 682 | 204 | 118.08 | + | 1.73 | 1.45E-11 | 1.17E-09 |
| signal transduction (GO:0007165) | 1136 | 298 | 196.68 | + | 1.52 | 7.61E-11 | 4.73E-09 |
| regulation of immune system process (GO:0002682) | 127 | 64 | 21.99 | + | 2.91 | 7.87E-11 | 4.75E-09 |
| leukocyte activation (GO:0045321) | 57 | 40 | 9.87 | + | 4.05 | 2.04E-10 | 1.11E-08 |
| positive regulation of immune system process (GO:0002684) | 87 | 50 | 15.06 | + | 3.32 | 2.25E-10 | 1.19E-08 |

### Proinflammatory Macrophages Increase Endometriotic Organoid Invasion

Immune cell dysfunction and local inflammation in the endometriotic environment have been considered to play a pivotal role in the pathogenesis of endometriosis (*20*). The recruitment of macrophages within the endometriotic lesion have been demonstrated to facilitate the development and maintenance of endometriosis (*21-23*). In support of a role for macrophages in the pathophysiology of endometriosis, immunoreactive CD68, a pan-macrophage marker, was detected in a baboon endometriotic lesion (Supplemental Fig 3). Proinflammatory, or M1, macrophages have the capacity to secrete inflammatory cytokines, including IL-6. These inflammatory cytokines may further promote endometriotic cell growth and invasion (*24*). To address the effect of activated proinflammatory macrophages on EO invasion, M1 were differentiated from naïve (M0) macrophages derived from the THP1 cell line by exposure to IFN-γ and lipopolysaccharide (LPS) for 24 hours. The resulting proinflammatory macrophages were acclimatized for 24-36 hours in the organoid culture medium. As before, EO added to the top of the Matrigel/LP9 layer and proinflammatory macrophages were added to the wells three hours later. The EO were imaged on days 1 and 3 using fluorescent laser-scanning confocal microscopy. Co-culture with either 7,500 or 15,000 macrophages increased EO invasion by 1.6-or 1.7-fold, respectively, while there was no difference between the macrophage treatment groups (Fig 7). Thus, the addition of proinflammatory macrophages promoted the endometriotic organoid invasion in support of the hypothesis that macrophages facilitate the establishment of endometriotic lesions.

**Fig 7.**
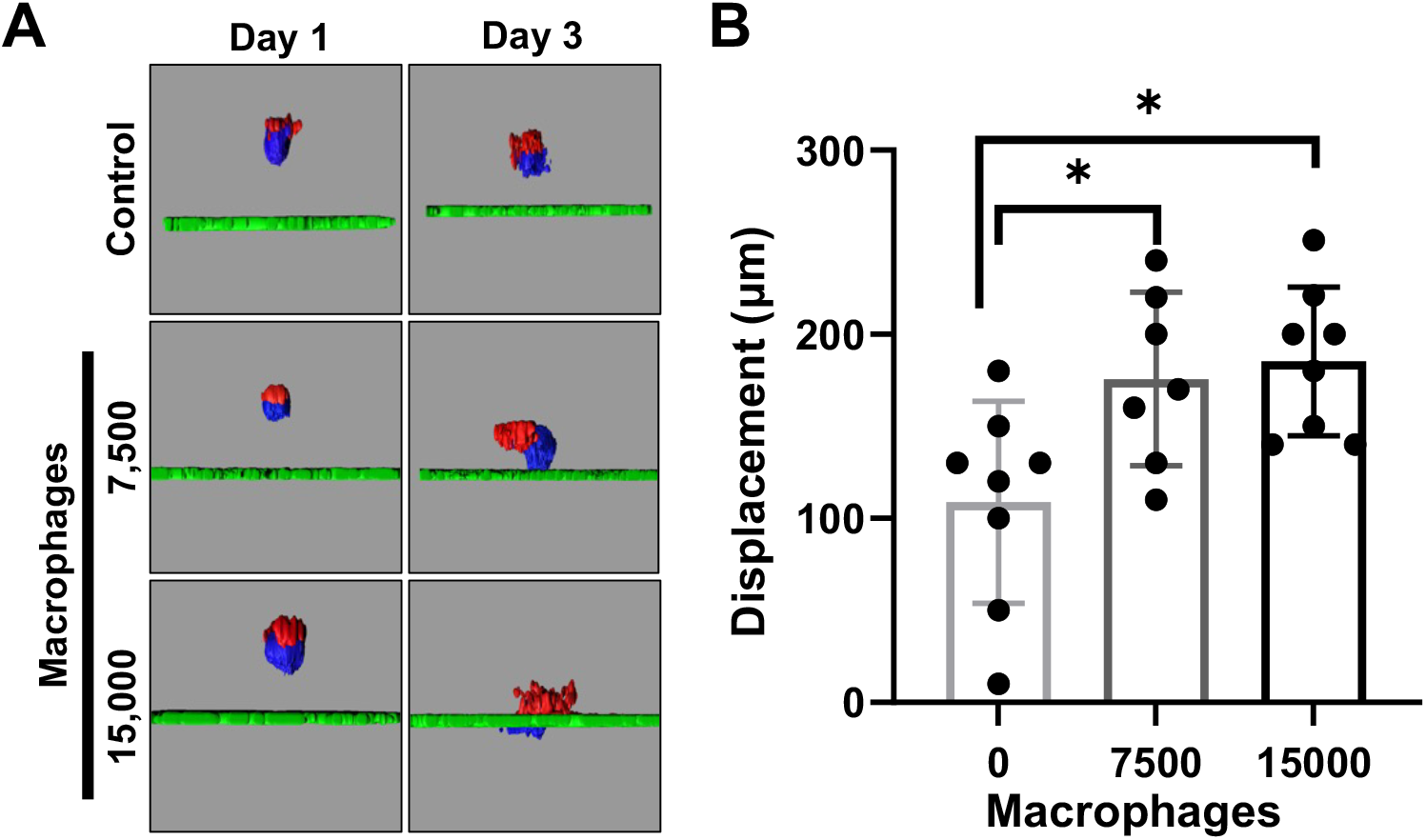
Proinflammatory macrophages increase EO invasion. (A) Day 4 EO were seeded on an LP9/Matrigel layer and M1 macrophages (7,500 or 15,000) were added to the plates after 3h. The EO were imaged on days 1 and 3 using fluorescent confocal microscopy and (B) invasion relative to the LP9 cell layer, was quantified. Invasion of EO was increased in co-culture with 7,500 or 15,000 macrophages by 1.6-or 1.7-fold, respectively. Control n=8; M1 7,500 n=7; M1 15,000 n=8 *P<0.05

## DISCUSSION

A 3D organoid consisting of both epithelial and stromal cells hold promise as an excellent model for dissecting mechanisms associated with early endometriotic lesion development. Most cases of endometriosis are hypothesized to be the result of retrograde menstruation of endometrial fragments containing both epithelium and stroma (*25, 26*). Therefore, inclusion of both cell types more closely resembles the pathophysiology of endometriosis and advances the potential for *in vitro* discovery. Ultra-low attachment plates and hanging drop culture have been used to generate spheroids with 12Z cells, immortalized stromal cells or primary ectopic stromal cells, or a combination of epithelial and stromal cells to investigate endometriosis (*27, 28*). Interestingly, both of these culture models formed self-organizing spheroid structures, but separate epithelial and stromal cell compartments were evident only in ultra-low attachment conditions (*27*); indicating that ultra-low attachment culture might be more suitable for developing organoids with multiple cell types.

In this study, we developed EO with endometriotic epithelial and stromal cell lines using a micro-molded agarose plate culture system. Organoids self-organized and resulted in a stromal cell core surrounded by epithelial cells, a morphology is similar to that of endometrial organoids derived from primary endometrial cells (*13*). Interestingly, the origin of the stromal cells, normal or ectopic, altered the organoid structure with stratified epithelial cells forming only with ectopic stromal cells. The observed epithelial stratification is consistent with reports of endometriotic epithelial organoids grown in Matrigel (*14*) and suggests that the ectopic stromal cells promoted epithelial cell proliferation and loss of apical-basal polarity. Transcriptomic analyses revealed that known pathways dysregulated in endometriotic lesions were present in EO, supporting the idea that EO reproduce many hallmarks of the endometriosis-associated transcriptome. Of note, normal endometrial epithelial cell lines were not included in this study and are not currently readily available, although an SV40-transfromed endometrial epithelial cell line, hEM3 was recently reported (*29*). Therefore, some of the gene signatures obtained from the classic comparison of endometriotic lesion versus normal endometrium may be absent from our study due to the limitation that both organoid types included diseased epithelium. This comparison, however, highlighted the effect of stromal cell origin on epithelial phenotype and poses important questions regarding the contribution of each cell compartment to the pathophysiology of lesions. Overall, our results support the idea that EO largely recapitulate the properties of endometriotic lesions with respect to morphology, function, and gene expression.

Attachment and invasion are important steps in the development of endometriotic lesions (*4*). Most current methods to study cell invasion are based on single cell types in 2D culture or Transwell assays. We developed a 3D organoid invasion model based on peritoneal endometriotic lesions to more accurately simulate the cell invasion process that occurs *in vivo*. To our knowledge, this is the first 3D invasion model which includes the three main cell compartments, epithelium, stroma and peritoneal mesothelium. Endometriotic organoids migrated through a layer of Matrigel, contacted, and penetrated through a layer of mesothelial cells, to mimic peritoneal invasion. Interestingly, our results identified stromal cells at the leading edge of invasion despite their location at the center of the organoid. This observation agrees with previous studies investigating adhesion and invasion of endometrium to peritoneum using both nude mice and *ex vivo* tissue explant culture models (*30, 31*). Endometrial stromal cells from women with endometriosis undergo changes in adhesive properties and mesenchymal characteristics in response to peritoneal mesothelial cells that might be part of the underlying mechanisms of lesion initiation (*32*). Our data indicate that endometriotic organoids have the capacity to invade a monolayer of mesothelial cells in a manner hypothesized to occur in physiological conditions. The utility of this model was further supported by increased invasive capacity in the presence of estrogen or activated proinflammatory macrophages. Collectively, this model expands the possibilities to study mechanisms of endometriotic lesion establishment *in vitro*.

Stromal cell origin significantly altered the transcriptome of endometriotic organoids with over 4,500 DEG. These observations support the utility of a 3D organoid comprised of endometriotic epithelium and stroma. Indeed, the genes expressed by EO largely mirrored those from spontaneous endometriotic lesions in the baboon, and GSEA analyses confirmed coordinated increases in TNF*α* signaling via NFKB, inflammatory responses, and hypoxia associated gene sets. Analysis of organoid gene expression alterations associated with the origin of the stromal cells identified several pathways known to be altered in endometriotic lesions, for example, retinoic acid (RA) signaling, both the uptake of retinoid (*33*) and the production of retinoic acid (RA) (*34*). Several enzymes in the RA pathway were altered in endometriotic organoids, including decreased expression of *STRA6* and *CRABP2* (*35*), and increased *CYP26B1*. Telomerase reverse transcriptase (*TERT*) expression was increased in ectopic organoids, as *TERT* was amongst the most highly upregulated genes but was not altered in a comparison of stroma-only organoids (Supplemental Fig 2). There is significant evidence for increased expression of *TERT* in endometriosis in eutopic secretory endometrium (*36-40*) and endometriotic lesions (*41*). In further support of the model, endometriotic lesions are progestin resistant (*42*) and, concordantly, progesterone receptor expression was downregulated in EO. The expression of leukemia inhibitory factor (*LIF*) was increased in EO, possibly due to local estrogenic activity (*43*) or inflammatory signals (*44, 45*). These results demonstrate significant alignment of the EO transcriptome with spontaneous disease and specific dysregulated pathways in endometriosis.

The immune system plays an important role in the establishment and progression of endometriosis (*24, 46, 47*). Tissue resident immune cells are refluxed with menstrual tissue and additional cells are recruited into the peritoneum where they interact with endometriotic lesions (*48*). The pro-inflammatory cytokine *IL6* is common to all three coordinately increased hallmark gene sets from EO and spontaneous endometriotic lesions from baboons (data not shown). Interleukin-6 expression was increased in the spontaneous baboon lesions and EO and is associated with chronic inflammation, endometriosis, and the presence of macrophages (*21, 48-50*). IL-6 activates the expression of *CCL2* (also referred to as monocyte chemoattractant protein 1 or *MCP1*) in endothelial cells to recruit monocytes (*51*) and IL-6 is further produced by the recruited macrophages and dendritic cells. Indeed, macrophages affect lesion establishment and growth in a mouse model of endometriosis (*21, 47, 48*) and peritoneal macrophages from women with endometriosis produce more IL-6 (*52*). Genes expressed in spontaneous lesions, but absent from EO, were enriched in immune-related pathways indicating a role for immune cells in the pathophysiology of endometriotic lesions that was absent in the model. Importantly, the invasive capacity of EO was increased with the addition of proinflammatory macrophages, further supporting this hypothesis. The mechanism by which the macrophages altered EO invasion is undetermined, but IL-6 may be a key cytokine for the crosstalk between endometriotic lesions and infiltrating macrophages. Our organoid model is an ideal system to uncover the crosstalk between lesions and immune cells and is a subject for future studies.

Here we describe scaffold-free self-organized endometriotic organoids, comprised of both endometriotic epithelial and stromal cells, combined with a novel model of attachment and invasion of the peritoneum. Our findings support the idea that 3D culture systems allow for the study of complex cell interactions and pathologically relevant processes like invasion and are a promising new pre-clinical tool to evaluate therapeutic treatments for endometriosis. EO recapitulated many of the transcriptomic features of spontaneous peritoneal endometriotic lesions from the baboon and the model allowed for real-time imaging and quantification of organoid invasion. The complexity of this model could be increased by altering conditions with exogenous compounds or other cell types. Future studies utilizing this system can capitalize on this flexibility with combinations of additional cell types, cell type genomic editing, and pharmacological compounds. At present our *in vitro* model does not reflect the full complexities of macrophage populations observed *in vivo* (*21, 47, 48*) and future experiments should investigate the interaction of peripheral monocytes or naïve macrophages on EO and determine the effect on all cell types, including the resultant macrophage phenotypes. Furthermore, EO could be integrated with microfluidics cultures containing endometrium (*53, 54*) to study endometriosis associated endometrial receptivity failure and decidualization defects.

In conclusion, our study suggests that EO recapitulate the characteristics of endometriotic lesions and support the concept that organoids are an appropriate model for dissecting the mechanisms that contribute to endometriotic lesion establishment.

## MATERIALS AND METHODS

### Cell Lines

Immortalized human endometriotic epithelial cells (12Z cells) (*16*) were cultured using Dulbecco’s modified Eagle’s medium (DMEM)/F-12 (Gibco, USA) supplemented with 10% Charcoal-dextran treated fetal bovine serum (CDS-FBS; Gibco, USA), 1X Pen/Strep (Gibco, USA) and 1X sodium pyruvate (Gibco, USA)(*16, 55*). The human peritoneal mesothelial cell line LP9 was kindly provided by Dr. James G. Rheinwald (Harvard Medical School, Boston, MA, USA). LP9 cells were maintained in a 1:1 ratio of M199 and Ham F12 media (Gibco), supplemented with 15% fetal bovine serum, 1% Penicillin/Streptomycin, 1% HEPES (Gibco), 1% GlutaMAX (Gibco), 10 ng/ml epidermal growth factor (EGF; Sigma, St Louis, MO, USA) and 400 ng/ml hydrocortisone (Sigma). The immortalized human endometriotic stromal cell line (iEc-ESC) was established in our laboratory (*17*). iEc-ESCs were cultured with phenol red–free DMEM/F12 medium supplemented with 10% charcoal dextran stripped fetal bovine serum (CDS-FBS), 100 units/ml of penicillin/streptomycin, and 1 mM sodium pyruvate. An immortalized human uterine fibroblast cell line (iHUF) was established from the decidua parietalis from term placenta in our laboratory. iHUF cells were cultured in phenol-red-free RPMI 1640 (Thermo Fisher Scientific) supplemented with 10% CDS-FBS, 100 units/ml of penicillin/streptomycin, and 1 mM sodium pyruvate. THP1 cells (kindly provided by Dr. Margaret Petroff, Michigan State University) were grown in T75 flask using RPMI1640 with 10% FBS in the presence of Antibiotic-Antimycotic in the suspension culture. These cells were sub-cultured either in 6-well plates or T-25 flask and treated with 100nM (1:5000 µl of media) PMA (Phorbo 12-Myristate 13-Acetate, Cayman Chemicals, USA) to yield 60-70% adherent cells at the end of 24 hours. The PMA exposure was repeated, and any floating non-confluent cells were removed. The cell obtained following PMA treatment were designated as M0 macrophages and kept in pre-differentiated (M0) state for 2-4 days before exposing them to M1/ M2 differentiation. These M0 adherent macrophages were activated with 20 ng/ml IFN-gamma (R&D Systems, USA) and 10pg/ ml of LPS (Sigma, USA.) for 24 hours to differentiate into M1 proinflammatory macrophages utilized to study effect of proinflammatory M1 macrophage on endometriotic organoid culture system. All cell lines were maintained at 37°C in an atmosphere of 5% CO_2_ in air.

RFP lentiviral vector (Ploc-RFP) was used to generate fluorescently tagged 12Z-RFP cell line. EGFP lentiviral vector (pHX-EGFP) was utilized to generate LP9-GFP stable cell lines. iEC-ESCs and iHUF cells were tagged with Azurite blue using pLV-Azurite (Addgene plasmid 36086). Lentiviral transductions were performed according to manufacturers’ protocols and successfully tagged cells were further sorted on a Beckman Coulter MoFlow Astrios (Indianapolis, IN, USA).

### Generation of Endometriotic Organoids

12Z and stromal cells were trypsinized in the flask and cell suspensions were centrifuged at 200 x*g* for 5 minutes. Both epithelial and stromal cell pellets were then suspended in complete MammoCult human medium (STEMCELL Technologies, Inc., 05620) supplemented with hydrocortisone and heparin as per the manufacturer’s instructions and 1% pen/strep (Sigma, P0781) to similar densities. Epithelial and stromal cells were mixed at a 1:50 ratio by volume and 50uL of epithelial-stromal cell suspension was seeded into 1.5% (w/v) agarose 3D micro-molded plate, prepared according to the manufacturer’s instructions (MicroTissues® 3D Petri Dish® micro-mold spheroids, Sigma, Z764043). Media was changed every 2-3 days. 12Z+iHUF organoids were used as a control, since a normal epithelial cell line of endometrial origin is not currently available.

### Organoid Invasion Model

To mimic the structure of the peritoneum, an ice cold LP9 cell suspension was mixed with ice cold Matrigel (Corning, 536232) at a ratio of 2:1(v:v), 30ul drops of Matrigel-cell suspension were plated into 96-well plate, at a density of 2-2.5×10^4^ cells per well, allowed to set at 37°C for 3-4 hours until solidified and overlaid with 100ul MammoCult growth medium. The next day, the day 4 endometriotic organoids were harvested from 3D Petri Dishes® by pipetting and then seeded on to the top of Matrigel-LP9-GFP layer in 96-well plates in 100ul MammoCult growth medium. Organoid invasion into the LP9-GFP/Matrigel layer were imaged every 48 hours using confocal microscopy.

### Organoid Imaging

Organoids were imaged using a Nikon Eclipse Ti inverted microscope using a Nikon C2 + confocal microscope laser scanner. For each organoid, z-stacks were generated with a 10μm increment, and tiled scans were set up to image the entire width and depth of the organoid with 10x air objective. The position of the each organoid was recorded as x,y coordinate by ND Acquisition of NIS-Elements software. For organoid sections, z-stacks were generated with 0.5μm increment using 40X oil objective. Images were merged using NIS-Elements C2 version 4.13.

### Quantification of Invasion

Commercial software Imaris x64 7.4.2 (Bitplane) was used for image analysis. In order to quantify the invasion of the organoid stromal cells into the peritoneal mesothelial cells, 3D surface models of the LP9-GFP cell layer, and the stromal cells-Azurite blue were created. Using the measurement function in Imaris, and using orthogonal slicers in XY, YZ and ZX planes, the linear distance between the organoid tip and the organoid migrating front on to the peritoneal cells were calculated. The distances were calculated at different time points in order to calculate velocity of organoid movement as a function of time. Seven to ten organoids for each time point in different groups were evaluated to account for heterogeneity in organoid structure as well as movement.

### Immunofluorescence Staining

Agarose 3D Petri Dishes® containing organoids were removed from the tissue culture plate and the medium inside was removed gently by pipetting under a dissecting microscope. Organoids were sealed within the agarose dish with lukewarm 1.5% (w/v) agarose in 1×PBS. The sealed 3D Petri Dishes® containing organoids were then fixed in 4% paraformaldehyde in 1×PBS at 4°C overnight, rinsed briefly in 1×PBS, stored in 70% EtOH and processed for standard paraffin embedding and sectioning at 6μm. Organoid sections were dewaxed, and then rehydrated with a graded alcohol series, followed by heat-mediated antigen retrieval in citrate buffer (Vector Laboratories, H3300) and then hydrogen peroxide treatment. Sections were blocked for 1 h in 10% normal horse serum (Vector Laboratories, S-2000) in PBS and incubated overnight at 4°C in primary antibodies. Organoid sections were incubated in respective species specific fluorochrome-conjugated secondary antibodies following an overnight incubation with the primary antibodies. Slides were then mounted with Antifade Mounting Medium with DAPI (Vector Laboratories, H-1200). Images were captured with a Nikon Eclipse Ti inverted microscope using software from NIS Elements, Inc. (Nikon, Melville, NY). The primary antibodies used were Vimentin polyclonal antibody (5741S) from Cell Signaling Technology, Inc. (Waltham, MA). Cytokeratin, pan (mixture) monoclonal antibody (C2562), from Sigma-Aldrich (St. Louis, MO). The secondary antibodies were: DyLight 594 goat anti-rabbit IgG antibody (DI-1594) from Vector Laboratories (Burlingame, CA). Alexa Fluor® 488 AffiniPure donkey anti-mouse IgG (715-545-150) from Jackson ImmunoResearch Laboratories Inc. (West Grove, PA).

### RNA-sequencing and Data Analysis

For RNA-sequencing, the Arcturus PicoPure RNA Isolation Kit (Themo Fisher Scientific, 12204-01), including an on-column DNA digestion using the RNAse-free DNAse set (Qiagen; 79254), was used to purify RNA from whole organoids. Total RNA was isolated from snap-frozen tissues homogenized in TRIzol reagent (Thermo Fisher; 15596018) following the manufacturer’s instructions. RNA was stored at -80°C in nuclease-free water and concentration was determined with a Nanodrop 1000 instrument (Thermo Fisher Scientific). Samples were sent to a sequencing facility (Novogene Corporation Inc., Chula Vista, CA) for RNA integrity analysis, library preparation, and sequencing. Libraries were prepared with an TruSeq mRNA kit (Illumina Inc., San Diego, CA) and sequenced (paired end 150 bp) on a NovaSeq 6000 instrument (Illumina Inc.) to an average depth of 39 million fragments per sample. Reads were quality trimmed and adapters were removed using TrimGalore (version 0.6.5) (*56*), and quality trimmed reads were assessed with FastQC (version 0.11.7). Trimmed reads were mapped to *Homo sapiens* genome assembly GRCh38 (hg38) or *Papio anubis* (version 3.0) using using HISAT2 (version 2.0.3) (*57*). Reads overlapping Ensembl annotations (version 99) (*58*) were quantified with featureCounts (version 1.6.2) (*59*) prior to model-based differential expression (DE) analysis using the edgeR-robust method (version 4.0.3) (*60*) in R. Genes with low counts per million (CPM) were removed using the filterByExpr function from edgeR. Transcriptome-wide read distributions were plotted to ensure a similar distribution across all samples with boxplots of log-transformed CPM values. Multidimensional scaling plots, generated with the plotMDS function of edgeR, were used to verify group separation prior to statistical analysis. Principal component analysis of normalized counts was conducted with the prcomp function in R using the top 500 variable genes. Differentially expressed genes (DEG) were identified as false discovery rate (FDR) p-value less than 0.05.

The hclust package was used for hierarchical clustering (ward.D) of Euclidean distances. Heat maps were generated using log-transformed transcript per million values (TPM) in pheatmap (version 1.0.12). Where necessary, baboon Ensembl genes were converted to human entrez identifiers with the human ortholog table downloaded from Biomart (version 101). The phyper function in R was used for hypergeometric tests of overlapping genes. Gene set enrichment and over-representation analyses were completed with clusterProfiler package (version 3.16.1) using mSigDB and KEGG reference databases.

### Statistical Analysis

Unless otherwise stated, statistical analysis was performed using GraphPad Prism 9.00 (GraphPad). All data were expressed as mean ± SD. The student’s t test was used for comparisons between the 2 groups, and 1-way ANOVA with a post hoc test least-significant difference was used for multiple comparisons. P < 0.05 was considered statistically significant (2-tailed).

## LIST OF SUPPLEMENTARY MATERIALS

Fig. S1. Epithelial and stromal cell organization in organoid culture.

Fig. S2. Telomerase reverse transcriptase (TERT) expression in organoids.

Fig. S3. Macrophages present in baboon endometriotic lesions.

Movie S1. Endometriotic organoids self-organize into epithelium and stromal compartments.

Movie S2. Confocal live imaging of EO invasion through a mesothelial cell layer over 8 days.

## ACKNOWLEDGEMENTS

The authors would like to sincerely thank all members of the Fazleabas laboratory, particularly Samantha Hrbek, Erin Vegter, and Ariadna Ochoa-Bernal, for their contributions to this project. We also thank Stephanie Celano for her assistance with live-cell imaging.

## FUNDING

National Institutes of Health grant R01HD083273 (ATF)

National Institutes of Health grant R01HD099090 (ATF)

National Institutes of Health grant F32HD104478 (GWB)

Endometriosis Foundation of America (ATF)

## AUTHOR CONTRIBUTIONS

Conceptualization: YS, GWB, ATF

Methodology: RA, JK

Investigation: YS, GWB, NRJ, RA

Visualization: YS, RA

Funding acquisition: GB, ATF

Project administration: ATF

Writing – original draft: YS, GB

Writing – review & editing: YS, GB, ATF, RA, JK

## COMPETING INTERESTS

Authors declare that they have no competing interests.

## DATA AND MATERIALS AVAILABILITY

Raw FASTQ files were deposited in the NCBI Gene Expression Omnibus (GSE######).

## Notes

### Competing Interest Statement

The authors have declared no competing interest.

## REFERENCES

1. L. C. Giudice, Clinical practice. Endometriosis. N Engl J Med 362, 2389–2398 (2010).

2. Practice Committee of the American Society for Reproductive Medicine. Treatment of pelvic pain associated with endometriosis. Fertil Steril 90, S260–269 (2008).

3. Endometriosis and infertility: a committee opinion. Fertil Steril 98, 591–598 (2012).

4. R. O. Burney, L. C. Giudice, Pathogenesis and pathophysiology of endometriosis. Fertil Steril 98, 511–519 (2012).

5. H. L. Yang, W. J. Zhou, K. K. Chang, J. Mei, L. Q. Huang, M. Y. Wang, Y. Meng, S. Y. Ha, D. J. Li, M. Q. Li, The crosstalk between endometrial stromal cells and macrophages impairs cytotoxicity of NK cells in endometriosis by secreting IL-10 and TGF-β. Reproduction 154, 815–825 (2017).

6. A. T. Fazleabas, A baboon model for inducing endometriosis. Methods Mol Med 121, 95–99 (2006).

7. P. Harirchian, I. Gashaw, S. T. Lipskind, A. G. Braundmeier, J. M. Hastings, M. R. Olson, A. T. Fazleabas, Lesion kinetics in a non-human primate model of endometriosis. Hum Reprod 27, 2341–2351 (2012).

8. K. L. Bruner-Tran, S. Mokshagundam, J. L. Herington, T. Ding, K. G. Osteen, Rodent Models of Experimental Endometriosis: Identifying Mechanisms of Disease and Therapeutic Targets. Curr Womens Health Rev 14, 173–188 (2018).

9. F. Schutgens, M. C. Verhaar, M. B. Rookmaaker, Pluripotent stem cell-derived kidney organoids: An in vivo-like in vitro technology. Eur J Pharmacol 790, 12–20 (2016).

10. S. L. Leibel, R. N. McVicar, A. M. Winquist, W. D. Niles, E. Y. Snyder, Generation of Complete Multi-Cell Type Lung Organoids From Human Embryonic and Patient-Specific Induced Pluripotent Stem Cells for Infectious Disease Modeling and Therapeutics Validation. Curr Protoc Stem Cell Biol 54, e118 (2020).

11. M. Boretto, B. Cox, M. Noben, N. Hendriks, A. Fassbender, H. Roose, F. Amant, D. Timmerman, C. Tomassetti, A. Vanhie, C. Meuleman, M. Ferrante, H. Vankelecom, Development of organoids from mouse and human endometrium showing endometrial epithelium physiology and long-term expandability. Development 144, 1775–1786 (2017).

12. M. Y. Turco, L. Gardner, J. Hughes, T. Cindrova-Davies, M. J. Gomez, L. Farrell, M. Hollinshead, S. G. E. Marsh, J. J. Brosens, H. O. Critchley, B. D. Simons, M. Hemberger, B. K. Koo, A. Moffett, G. J. Burton, Long-term, hormone-responsive organoid cultures of human endometrium in a chemically defined medium. Nat Cell Biol 19, 568–577 (2017).

13. T. Wiwatpanit, A. R. Murphy, Z. Lu, M. Urbanek, J. E. Burdette, T. K. Woodruff, J. J. Kim, Scaffold-Free Endometrial Organoids Respond to Excess Androgens Associated With Polycystic Ovarian Syndrome. J Clin Endocrinol Metab 105, (2020).

14. M. Boretto, N. Maenhoudt, X. Luo, A. Hennes, B. Boeckx, B. Bui, R. Heremans, L. Perneel, H. Kobayashi, I. Van Zundert, H. Brems, B. Cox, M. Ferrante, I. H. Uji, K. P. Koh, T. D’Hooghe, A. Vanhie, I. Vergote, C. Meuleman, C. Tomassetti, D. Lambrechts, J. Vriens, D. Timmerman, H. Vankelecom, Patient-derived organoids from endometrial disease capture clinical heterogeneity and are amenable to drug screening. Nat Cell Biol 21, 1041–1051 (2019).

15. T. M. Rawlings, K. Makwana, D. M. Taylor, M.A. Molè, K. J. Fishwick, M. Tryfonos, J. Odendaal, A. Hawkes, M. Zernicka-Goetz, G. M. Hartshorne, J. J. Brosens, E. S. Lucas, Modelling the impact of decidual senescence on embryo implantation in human endometrial assembloids. Elife 10, (2021).

16. A. Zeitvogel, R. Baumann, A. Starzinski-Powitz, Identification of an invasive, N-cadherin-expressing epithelial cell type in endometriosis using a new cell culture model. Am J Pathol 159, 1839–1852 (2001).

17. Y. Song, N. R. Joshi, E. Vegter, S. Hrbek, B. A. Lessey, A. T. Fazleabas, Establishment of an Immortalized Endometriotic Stromal Cell Line from Human Ovarian Endometrioma. Reproductive sciences (Thousand Oaks, Calif.), (2020).

18. W. Xiong, L. Zhang, L. Yu, W. Xie, Y. Man, Y. Xiong, H. Liu, Y. Liu, Estradiol promotes cells invasion by activating β-catenin signaling pathway in endometriosis. Reproduction 150, 507–516 (2015).

19. Y. Du, Z. Zhang, W. Xiong, N. Li, H. Liu, H. He, Q. Li, Y. Liu, L. Zhang, Estradiol promotes EMT in endometriosis via MALAT1/miR200s sponge function. Reproduction 157, 179–188 (2019).

20. S. H. Ahn, S. P. Monsanto, C. Miller, S. S. Singh, R. Thomas, C. Tayade, Pathophysiology and Immune Dysfunction in Endometriosis. Biomed Res Int 2015, 795976 (2015).

21. M. Bacci, A. Capobianco, A. Monno, L. Cottone, F. Di Puppo, B. Camisa, M. Mariani, C. Brignole, M. Ponzoni, S. Ferrari, P. Panina-Bordignon, A. A. Manfredi, P. Rovere-Querini, Macrophages are alternatively activated in patients with endometriosis and required for growth and vascularization of lesions in a mouse model of disease. Am J Pathol 175, 547–556 (2009).

22. E. Haber, H. D. Danenberg, N. Koroukhov, R. Ron-El, G. Golomb, M. Schachter, Peritoneal macrophage depletion by liposomal bisphosphonate attenuates endometriosis in the rat model. Hum Reprod 24, 398–407 (2009).

23. P. Vercellini, P. Vigano, E. Somigliana, L. Fedele, Endometriosis: pathogenesis and treatment. Nat Rev Endocrinol 10, 261–275 (2014).

24. G. Izumi, K. Koga, M. Takamura, T. Makabe, E. Satake, A. Takeuchi, A. Taguchi, Y. Urata, T. Fujii, Y. Osuga, Involvement of immune cells in the pathogenesis of endometriosis. J Obstet Gynaecol Res 44, 191–198 (2018).

25. J. A. Sampson, Peritoneal endometriosis due to the menstrual dissemination of endometrial tissue into the peritoneal cavity. American Journal of Obstetrics & Gynecology 14, 422–469 (1927).

26. K. T. Zondervan, C. M. Becker, K. Koga, S. A. Missmer, R. N. Taylor, P. Vigano, Endometriosis. Nat Rev Dis Primers 4, 9 (2018).

27. J. R. H. Wendel, X. Wang, L. J. Smith, S. M. Hawkins, Three-Dimensional Biofabrication Models of Endometriosis and the Endometriotic Microenvironment. Biomedicines 8, (2020).

28. A. Stejskalova, V. Fincke, M. Nowak, Y. Schmidt, K. Borrmann, M. K. von Wahlde, S. D. Schafer, L. Kiesel, B. Greve, M. Gotte, Collagen I triggers directional migration, invasion and matrix remodeling of stroma cells in a 3D spheroid model of endometriosis. Sci Rep 11, 4115 (2021).

29. Y. Park, J. G. Jung, Z. C. Yu, R. Asaka, W. Shen, Y. Wang, W. H. Jung, A. Tomaszewski, G. Shimberg, Y. Chen, V. Parimi, S. Gaillard, I. M. Shih, T. L. Wang, A novel human endometrial epithelial cell line for modeling gynecological diseases and for drug screening. Lab Invest 101, 1505–1512 (2021).

30. C. A. Witz, I. A. Monotoya-Rodriguez, R. S. Schenken, Whole explants of peritoneum and endometrium: a novel model of the early endometriosis lesion. Fertil Steril 71, 56–60 (1999).

31. M. Nisolle, F. Casanas-Roux, J. Donnez, Early-stage endometriosis: adhesion and growth of human menstrual endometrium in nude mice. Fertil Steril 74, 306–312 (2000).

32. L. L. Lin, S. Makwana, M. Chen, C. M. Wang, L. H. Gillette, T. H. Huang, R. O. Burney, B. J. Nicholson, N. B. Kirma, Cellular junction and mesenchymal factors delineate an endometriosis-specific response of endometrial stromal cells to the mesothelium. Mol Cell Endocrinol 539, 111481 (2021).

33. M. E. Pavone, S. Reierstad, H. Sun, M. Milad, S. E. Bulun, Y. H. Cheng, Altered retinoid uptake and action contributes to cell survival in endometriosis. J Clin Endocrinol Metab 95, E300–309 (2010).

34. K. Pierzchalski, R. N. Taylor, C. Nezhat, J. W. Jones, J. L. Napoli, G. Yang, M. A. Kane, N. Sidell, Retinoic acid biosynthesis is impaired in human and murine endometriosis. Biol Reprod 91, 84 (2014).

35. M. E. Pavone, A. R. Grover, R. Confino, E. K. Pearson, S. Malpani, Y.-H. Cheng, A. Fazleabas, S. Bulun, Retinoic acid action is altered within endometrium of baboons affected with endometriosis. Journal of Endometriosis and Pelvic Pain Disorders, (2022).

36. C. M. Kim, Y. J. Oh, S. H. Cho, D. J. Chung, J. Y. Hwang, K. H. Park, D. J. Cho, Y. M. Choi, B. S. Lee, Increased telomerase activity and human telomerase reverse transcriptase mRNA expression in the endometrium of patients with endometriosis. Hum Reprod 22, 843–849 (2007).

37. D. K. Hapangama, M. A. Turner, J. A. Drury, S. Quenby, G. Saretzki, C. Martin-Ruiz, T. Von Zglinicki, Endometriosis is associated with aberrant endometrial expression of telomerase and increased telomere length. Hum Reprod 23, 1511–1519 (2008).

38. A. J. Valentijn, K. Palial, H. Al-Lamee, N. Tempest, J. Drury, T. Von Zglinicki, G. Saretzki, P. Murray, C. E. Gargett, D. K. Hapangama, SSEA-1 isolates human endometrial basal glandular epithelial cells: phenotypic and functional characterization and implications in the pathogenesis of endometriosis. Hum Reprod 28, 2695–2708 (2013).

39. F. A. Mafra, D. M. Christofolini, V. Cavalcanti, F. L. Vilarino, G.M. André, P. Kato, B. Bianco, C. P. Barbosa, Aberrant telomerase expression in the endometrium of infertile women with deep endometriosis. Arch Med Res 45, 31–35 (2014).

40. D. K. Hapangama, A. Kamal, G. Saretzki, Implications of telomeres and telomerase in endometrial pathology. Hum Reprod Update 23, 166–187 (2017).

41. D. K. Hapangama, M. A. Turner, J. Drury, L. Heathcote, Y. Afshar, P. A. Mavrogianis, A. T. Fazleabas, Aberrant expression of regulators of cell-fate found in eutopic endometrium is found in matched ectopic endometrium among women and in a baboon model of endometriosis. Hum Reprod 25, 2840–2850 (2010).

42. G. R. Attia, K. Zeitoun, D. Edwards, A. Johns, B. R. Carr, S. E. Bulun, Progesterone receptor isoform A but not B is expressed in endometriosis. J Clin Endocrinol Metab 85, 2897–2902 (2000).

43. S. E. Bulun, S. Yang, Z. Fang, B. Gurates, M. Tamura, S. Sebastian, Estrogen production and metabolism in endometriosis. Annals of the New York Academy of Sciences 955, 75-85; discussion 86-78, 396-406 (2002).

44. D. Knight, Leukaemia inhibitory factor (LIF): a cytokine of emerging importance in chronic airway inflammation. Pulm Pharmacol Ther 14, 169–176 (2001).

45. G. X. Rosario, C. L. Stewart, The Multifaceted Actions of Leukaemia Inhibitory Factor in Mediating Uterine Receptivity and Embryo Implantation. Am J Reprod Immunol 75, 246–255 (2016).

46. D. I. Lebovic, M. D. Mueller, R. N. Taylor, Immunobiology of endometriosis. Fertil Steril 75, 1–10 (2001).

47. C. Hogg, A. W. Horne, E. Greaves, Endometriosis-Associated Macrophages: Origin, Phenotype, and Function. Front Endocrinol (Lausanne) 11, 7 (2020).

48. C. Hogg, K. Panir, P. Dhami, M. Rosser, M. Mack, D. Soong, J. W. Pollard, S. J. Jenkins, A. W. Horne, E. Greaves, Macrophages inhibit and enhance endometriosis depending on their origin. Proc Natl Acad Sci U S A 118, (2021).

49. C. Gabay, Interleukin-6 and chronic inflammation. Arthritis Res Ther 8 Suppl 2, S3 (2006).

50. A. Capobianco, A. Monno, L. Cottone, M. A. Venneri, D. Biziato, F. Di Puppo, S. Ferrari, M. De Palma, A. A. Manfredi, P. Rovere-Querini, Proangiogenic Tie2(+) macrophages infiltrate human and murine endometriotic lesions and dictate their growth in a mouse model of the disease. Am J Pathol 179, 2651–2659 (2011).

51. M. Romano, M. Sironi, C. Toniatti, N. Polentarutti, P. Fruscella, P. Ghezzi, R. Faggioni, W. Luini, V. van Hinsbergh, S. Sozzani, F. Bussolino, V. Poli, G. Ciliberto, A. Mantovani, Role of IL-6 and its soluble receptor in induction of chemokines and leukocyte recruitment. Immunity 6, 315–325 (1997).

52. P. Montagna, S. Capellino, B. Villaggio, V. Remorgida, N. Ragni, M. Cutolo, S. Ferrero, Peritoneal fluid macrophages in endometriosis: correlation between the expression of estrogen receptors and inflammation. Fertil Steril 90, 156–164 (2008).

53. H. Campo, A. Murphy, S. Yildiz, T. Woodruff, I. Cervelló, J. J. Kim, Microphysiological Modeling of the Human Endometrium. Tissue Eng Part A 26, 759–768 (2020).

54. S. Xiao, J. R. Coppeta, H. B. Rogers, B. C. Isenberg, J. Zhu, S. A. Olalekan, K. E. McKinnon, D. Dokic, A. S. Rashedi, D. J. Haisenleder, S. S. Malpani, C. A. Arnold-Murray, K. Chen, M. Jiang, L. Bai, C. T. Nguyen, J. Zhang, M. M. Laronda, T. J. Hope, K. P. Maniar, M. E. Pavone, M. J. Avram, E. C. Sefton, S. Getsios, J. E. Burdette, J. J. Kim, J. T. Borenstein, T. K. Woodruff, A microfluidic culture model of the human reproductive tract and 28-day menstrual cycle. Nat Commun 8, 14584 (2017).

55. S. K. Banu, J. Lee, A. Starzinski-Powitz, J. A. Arosh, Gene expression profiles and functional characterization of human immortalized endometriotic epithelial and stromal cells. Fertil Steril 90, 972–987 (2008).

56. M. Martin, Cutadapt removes adapter sequences from high-throughput sequencing reads. EMBnet.journal 17, pp–10 (2011).

57. D. Kim, B. Langmead, S. L. Salzberg, HISAT: A fast spliced aligner with low memory requirements. Nat Methods 12, 357–360 (2015).

58. B. L. Aken, S. Ayling, D. Barrell, L. Clarke, V. Curwen, S. Fairley, J. Fernandez Banet, K. Billis, C. Garcia Giron, T. Hourlier, K. Howe, A. Kahari, F. Kokocinski, F. J. Martin, D. N. Murphy, R. Nag, M. Ruffier, M. Schuster, Y. A. Tang, J. H. Vogel, S. White, A. Zadissa, P. Flicek, S. M. Searle, The Ensembl gene annotation system. Database (Oxford) 2016, 1–19 (2016).

59. Y. Liao, G. K. Smyth, W. Shi, featureCounts: an efficient general purpose program for assigning sequence reads to genomic features. Bioinformatics 30, 923–930 (2014).

60. X. Zhou, H. Lindsay, M. D. Robinson, Robustly detecting differential expression in RNA sequencing data using observation weights. Nucleic Acids Res 42, e91 (2014).

