## Supplemental Materials for "Endometriotic Organoids: A Novel In Vitro Model of Endometriotic Lesion Development"

### SUPPLEMENTARY MATERIALS

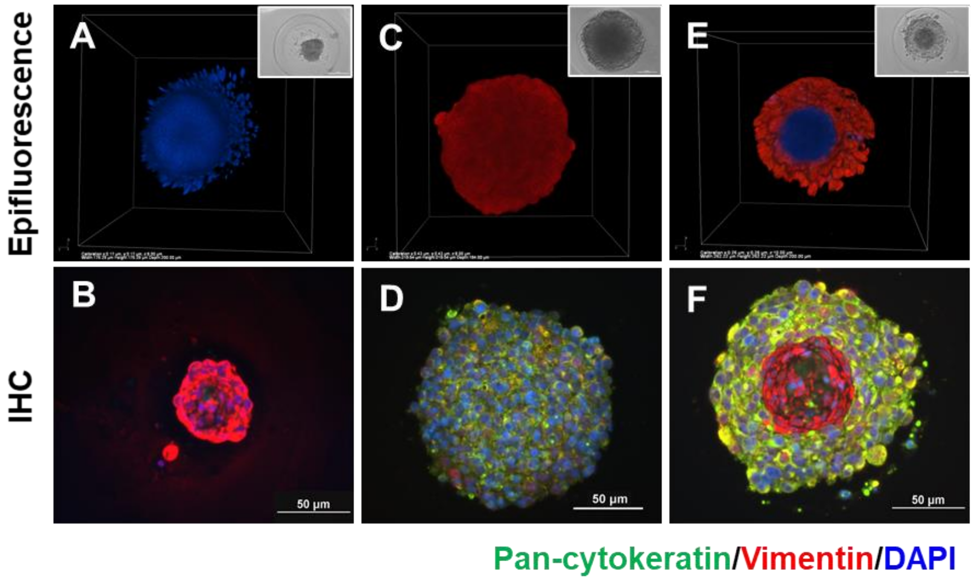


**Fig. S1. Epithelial and stromal cell organization in organoid culture.** Organoids made with (A-B) iEc-ESC-Azurite Blue alone, (C-D) 12Z-RFP alone, and (E-F) a combination of both cell types. (A,C,E) Epifluorescent 3D views of organoid structure. Inserts are matching phase contrast images. (B,D,F) Immunofluorescent organoid staining for vimentin (red) and cytokeratin (green).

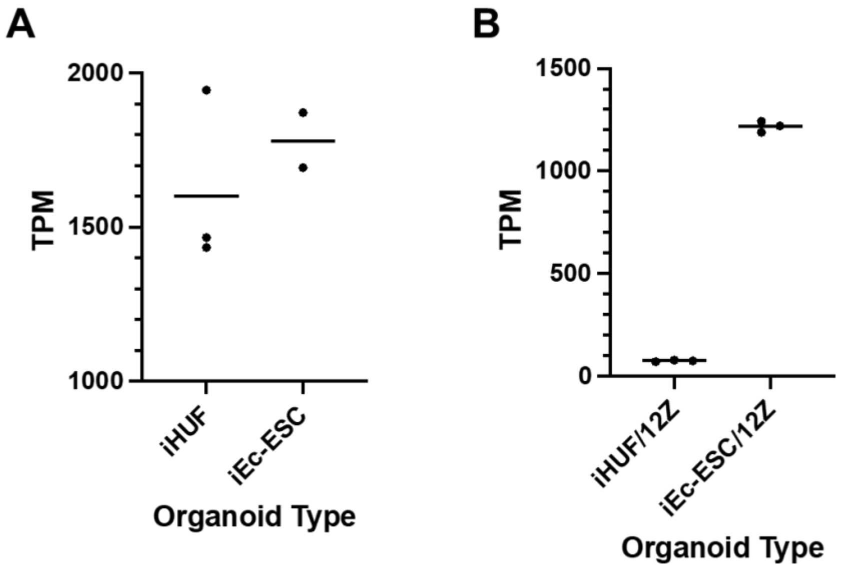


**Fig. S2. Telomerase reverse transcriptase (TERT) expression in organoids.** A) Expression was not different between stromal-cell only organoids (FDR p = 0.91). B) In contrast, TERT was one of the most highly increased genes in iEc-ESC/12Z organoids (FDR p = 1.3x10-307). Horizontal bars are the geometric mean.

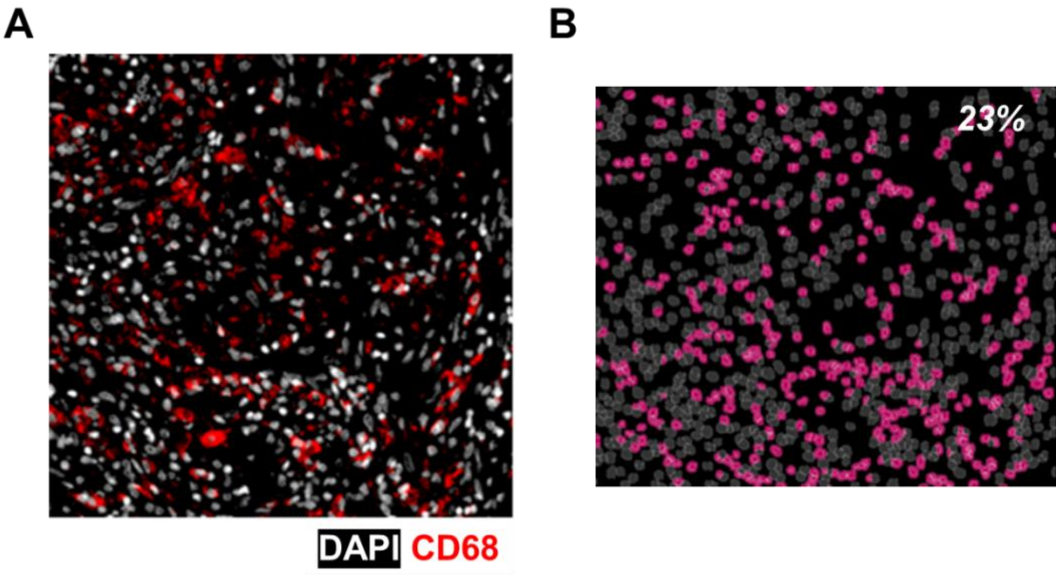


**Fig. S3. Macrophages are present in baboon endometriotic lesions.** A) Fluorescent immunohistochemical image of baboon endometriotic lesions stained with CD68 (red), a pan-macrophage marker. B) Quantification found 23% of cells were CD68 positive.

See separate video file.

**Movie S1. Endometriotic organoids self-organize into epithelium and stroma compartments.**

See separate video file.

**Movie S2. Confocal live imaging of EO invasion through a mesothelial cell layer over 8 days.**
